# Triphasic Thrombosis Model: A Computational Study of Type B Aortic Dissection

**DOI:** 10.1101/2024.05.07.592918

**Authors:** Ishan Gupta, Martin Schanz, Tim Ricken

## Abstract

Thrombosis refers to the formation of a thrombus, or a blood clot, within the body, which can occur either partially or completely. It serves as a crucial indicator of the severity of a patient’s medical condition, with the location and characteristics of thrombosis dictating its clinical implications. Hence, accurate diagnosis and effective management of thrombosis are paramount.

In our current investigation, we incorporate the porous attributes of a thrombus using the Theory of Porous Media. This involves dividing the aggregate into solid, liquid, and nutrient phases and utilising volume fractions to capture microstructural details. Fluid flow through the porous media is modelled using a modified Darcy-Brinkman type equation, with interaction terms within balance equations facilitating the modelling of the mass exchange and other phase interactions. The shorter time scales are neglected.

We present a comprehensive framework of equations and assumptions governing the behaviour of a strongly coupled multiphasic porous medium problem. Additionally, we introduce scenarios involving type B Aortic Dissection and false lumen geometries, providing a detailed outline of the problem setup. Thereafter, we present the potential of the model for thrombi growth. The simulation results are compared with velocity plots aligning with Magnetic Resonance Imaging data for three distinct cases with varying entry and exit tear sizes. Consequently, our proposed model offers a promising and reasonable approach for numerically simulating thrombosis and gaining insights into the underlying growth mechanics.

## 1 Introduction

### 1.1 Motivation

Thrombosis is the process of forming a spatial structure known as a thrombus (blood clot). The formation of a thrombus involves a complex sequence of biochemical reactions. While the body has a system to keep blood fluid and prevent bleeding, an injury to a blood vessel triggers haemostasis, which forms a thrombus to stop the bleeding. Thrombosis occurs in two main steps. First, platelets are activated and form a plug at the injury site. Second, the coagulation cascade is triggered, leading to the formation of fibrin fibres. The fibrin fibres, along with the platelets, create a stable, permanent plug called a thrombus.

Thrombosis can be partial or complete and is a crucial indicator of the severity of a medical condition. On the one hand, it can save a life by healing an injury, such as in the case of a skin laceration or an Aortic Dissection (AD). On the other hand, it can endanger life by blocking blood vessels, as seen in cases of venous or arterial thrombosis. Moreover, some patients can be at risk of both thrombosis and bleeding simultaneously, as in the case of Disseminated Intravascular Coagulation (DIC), which is a widespread hypercoagulable state. Thrombosis can occur in any part of the body, and the clinical consequences depend on the location and nature of the thrombosis. As a result, the diagnosis and management of thrombosis is crucial (Kumar et al. 2017; Mohan 2018). For this study, we choose the case of AD for the model development process. In this context, we will briefly introduce AD and the nature of thrombosis in this condition.

### 1.2 Aortic Dissection

Aorta is the largest blood vessel in the body and serves as the main artery carrying blood from the heart to the entire body. The heart pumps the blood from the left ventricle into the aorta via the aortic valve, which opens and closes with each heartbeat to allow a one-way blood flow. One of the most common forms of acute aortic syndrome is AD. It begins when a tear occurs in the inner layer (intima) of the aortic wall. This tear allows the blood to flow between the inner and middle layers causing them to separate (dissect). This second blood-filled channel is called the false lumen. This condition can lead to aortic rupture or decreased blood flow to the organs, causing short-or long-term damage. AD is a severe condition that can be fatal if not treated early. The risk and nature of AD complications are highly dependent on the affected area of the aorta. There are two types of AD depending on the location of the dissection, cf. Figure 1. In type A AD, dissection occurs in the ascending part of the aorta, where the expansion of the false lumen can compress other branches of the aorta and reduce blood flow. On the contrary, dissection occurs in the descending part of the aorta in type B AD, which may extend to the abdomen. Complete or partial thrombosis in the false lumen is a significant predictor of mortality in patients (Kumar et al. 2017; Tsai et al. 2007).

**Fig. 1.**
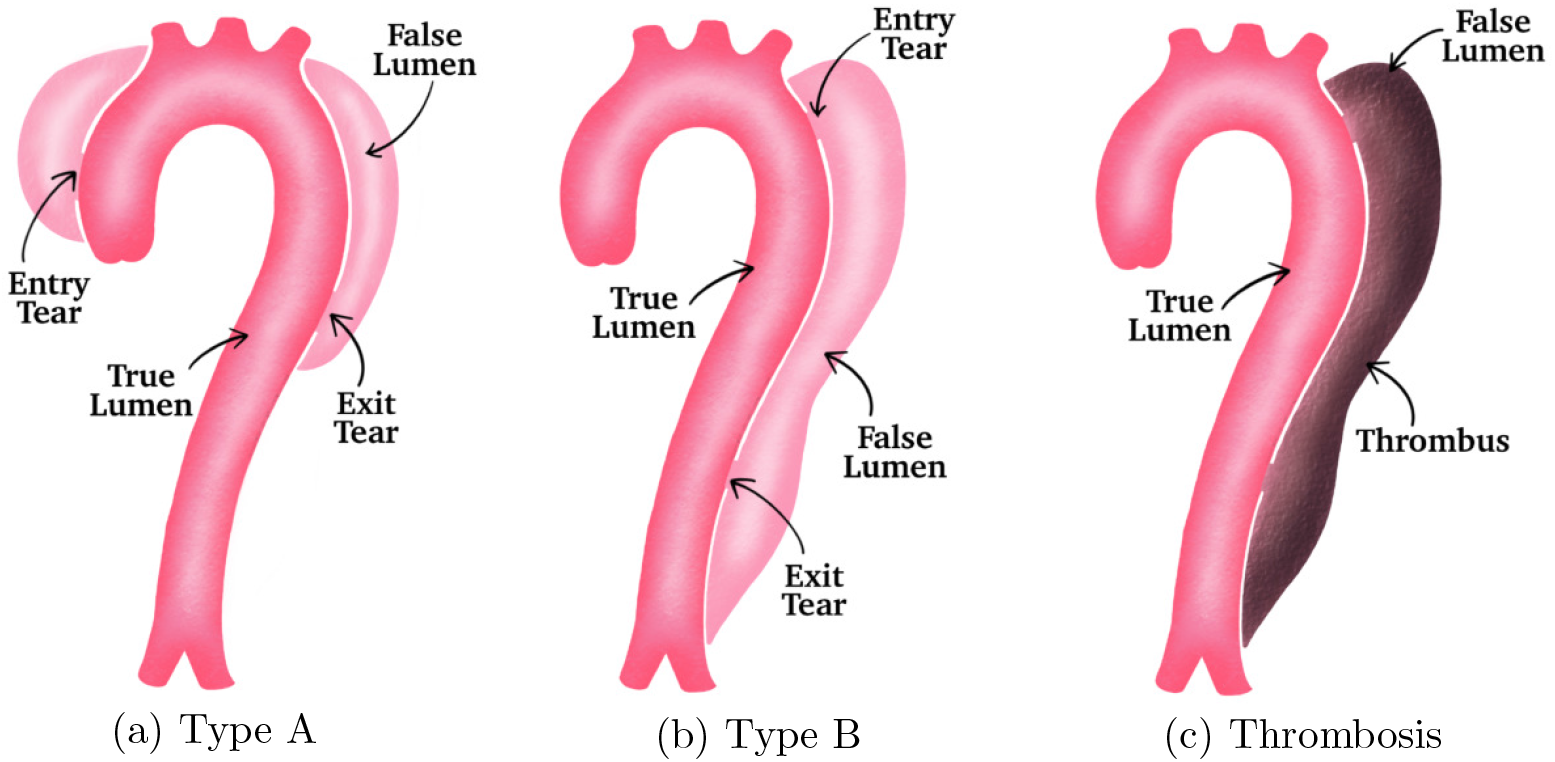
Illustrations of the entry tear, exit tear, true lumen, and false lumen in (a) type A AD, (b) type B AD, and (c) formation of thrombi in type B AD (Gupta 2023; Tsai et al. 2007)

AD is a severe condition that may arise due to various risk factors. High blood pressure leading to increased stress on the aortic wall is the most common risk factor. Other factors include weakening of the wall, pre-existing aneurysm or defects in the aortic valve. AD primarily affects two epidemiological groups. The first group is men aged 40 to 60 years with pre-existing hypertension, and the second is younger patients with systemic or localised connective tissue abnormalities affecting the aorta. The estimated occurrence of AD is 3 to 6 cases per 100,000 people annually in Europe and the United States (Hibino et al. 2022). Moreover, in the instance of type B AD, the mortality rate is excessively high in the first seven days due to severe complications, such as malperfusion or rupture in the aorta. Approximately 40% of the patients die immediately from AD. Moreover, only 50% to 70% live 5 years after the surgery depending on the age and underlying complications (Kumar and Allain, 2012). This has boosted the importance of early diagnosis and treatment, which is critical for survival. Even though a general agreement exists on the need for immediate surgical repair for patients with acute type A AD, the optimal treatment of type B AD remains a matter of debate (Terzi et al. 2018; Luebke and Brunkwall 2014; Fann et al. 1995). This led to an interest in computational methods to help with the decision-making process for treatment. Our current understanding of optimal medical care can expand by increasing insights into the progression and nature of the disease. A model for thrombosis prediction in patient-specific cases can help doctors make informed decisions and analyse the dangers involved.

### 1.3 State of the Art

The mathematical modelling of thrombosis first emerged in the late 1980s by Khanin and Semenov (1989). This research focused mainly on the coagulation network, which is a part of secondary haemostasis. Since then, the models have improved considerably. A cell-based model for haemostasis was presented by Hoffman and Monroe III (2001) which was a set of Ordinary Differential Equations (ODEs). This model was further enhanced by Lobanov et al. (2011). Ovanesov et al. (2002) introduced a reaction-diffusion model based on Partial Differential Equations (PDEs), which was further developed using Computational Fluid Dynamics (CFD) with increased complexity by Leiderman and Fogelson (2011), involving more than 50 PDEs. Chen and Diamond (2019) reduced the complexity of the model by using 8 ODEs, although these excluded platelet aggregation. Anand et al. (2022) provide a detailed review of haemostasis models.

Moreover, various methodologies were employed to develop models in the context of thrombosis. Storti and van de Vosse (2014) introduced a coupled solid mechanics and Fluid-Structure Interaction (FSI) model focusing on platelet plug formation. Chen et al. (2016) presented another model based on FSI with simplified geometry, capable of predicting blood flow velocity and volumes of true and false lumen. Cheng et al. (2013) introduced a CFD model for blood flow, treating blood as a Newtonian and incompressible fluid, specifically for the case of AD. Biasetti et al. (2010) developed a CFD model for thrombi formation mechanisms in abdominal aortic aneurysms. de Sousa et al. 2015 and Malaspinas et al. (2016) focused on the impact of haemodynamic parameters on initial thrombi formation using CFD models. Xu et al. (2008) presented a 2D multiscale model for thrombi development based on CFD. Menichini and Xu (2016) proposed a CFD model with mass transfer equations and idealised geometries, demonstrating good agreement with clinical data. Jafarinia et al. (2022) presented a shear-driven model for predicting thrombosis. However, the correlation between the simulation time and the physiological time remains a challenge. Armour et al. (2022) explored the haemodynamic impact of branches originating from the false lumen in cases of false lumen thrombosis. Additionally, Wang et al. (2023) emphasised the importance of incorporating thrombus breakdown in computational models of thrombosis under steady flow conditions. Despite certain limitations of these models, they do validate the applicability of computational models in thrombosis prediction.

Modelling a thrombus is a continuum-mechanical problem that cannot be uniquely classified within well-known solid or fluid mechanics disciplines. Instead, it is necessary to consider a multiphasic aggregate along with the associated characteristics of its constituents. During thrombosis, evolving tissues undergo changes in their internal structure and properties other than their shape and size. As platelets cohere into a platelet aggregate, they do not immediately form a solid mass. Instead, they create a porous network with voids and channels. Following numerous reactions, the thrombus undergoes a transition to a stiff solid state. The porous nature of a thrombus makes the use of a macroscopic continuum-mechanical approach like the Theory of Porous Media (TPM) more suitable for modelling. It provides an excellent framework to describe the overall behaviour of the multiphasic thrombus.

Various models based on TPM have been proposed to characterise the mechanical behaviour, growth, and remodelling processes of biological tissues. A biphasic model was proposed by Ehlers and Markert (2001) for describing the mechanical behaviour of hydrated soft tissues and by Wagner and Ehlers (2008) for describing the behaviour of brain tissue. Furthermore, Karajan (2012) introduced a biphasic model to describe the mechanical behaviour of the Intervertebral Disc (IVD). Ricken et al. (2010) presented a biphasic model for liver perfusion remodelling. Moreover, multiphasic models for growth and remodelling have been proposed by Ricken and Bluhm (2009, 2010) for isotropic and transversely isotropic biological tissues. Preziosi and Tosin (2009) presented multiphasic tumour growth models. Krause (2014) proposed a model for tumour growth and bone remodelling using the growth energy concept. The application of TPM to describe the behaviour and growth of soft biological tissues has been well summarised by Ateshian and Humphrey (2012). The recent works employing TPM for modelling biological growth further motivate its use and highlight its relevance for modelling thrombosis.

In a recent study, we introduced a multiphasic model for thrombosis using both idealised and realistic 2D geometries (Gupta and Schanz, 2023a,b; Gupta, 2023). However, in this investigation, we improve and extend the model to incorporate the viscous properties of the fluid, which become significant at high velocities. Additionally, we use actual 3D geometries to investigate the case of type B Aortic Dissection. Furthermore, we validate the simulation outcomes by comparing them with MRI scans. Additionally, we assess the model’s performance across various dimensions of the entry and exit tears of the false lumen. Nonetheless, it is crucial to recognise that modelling the growth of living tissues poses inherent challenges, as well summarised by Ambrosi et al. (2019).

## 2 Materials and Methods

### 2.1 The Theory of Porous Media

Within the developed framework of the Theory of Porous Media, we can obtain a macro-model to describe the complicated microstructure by using the homogenisation procedure over a Representative Elementary Volume (REV) of a thrombus, cf. Figure 2. We can split the whole constituent *φ* into three phases *φ*^*α*^, where *α* = {*S, L, N* }, leading to a triphasic model

**Fig. 2.**
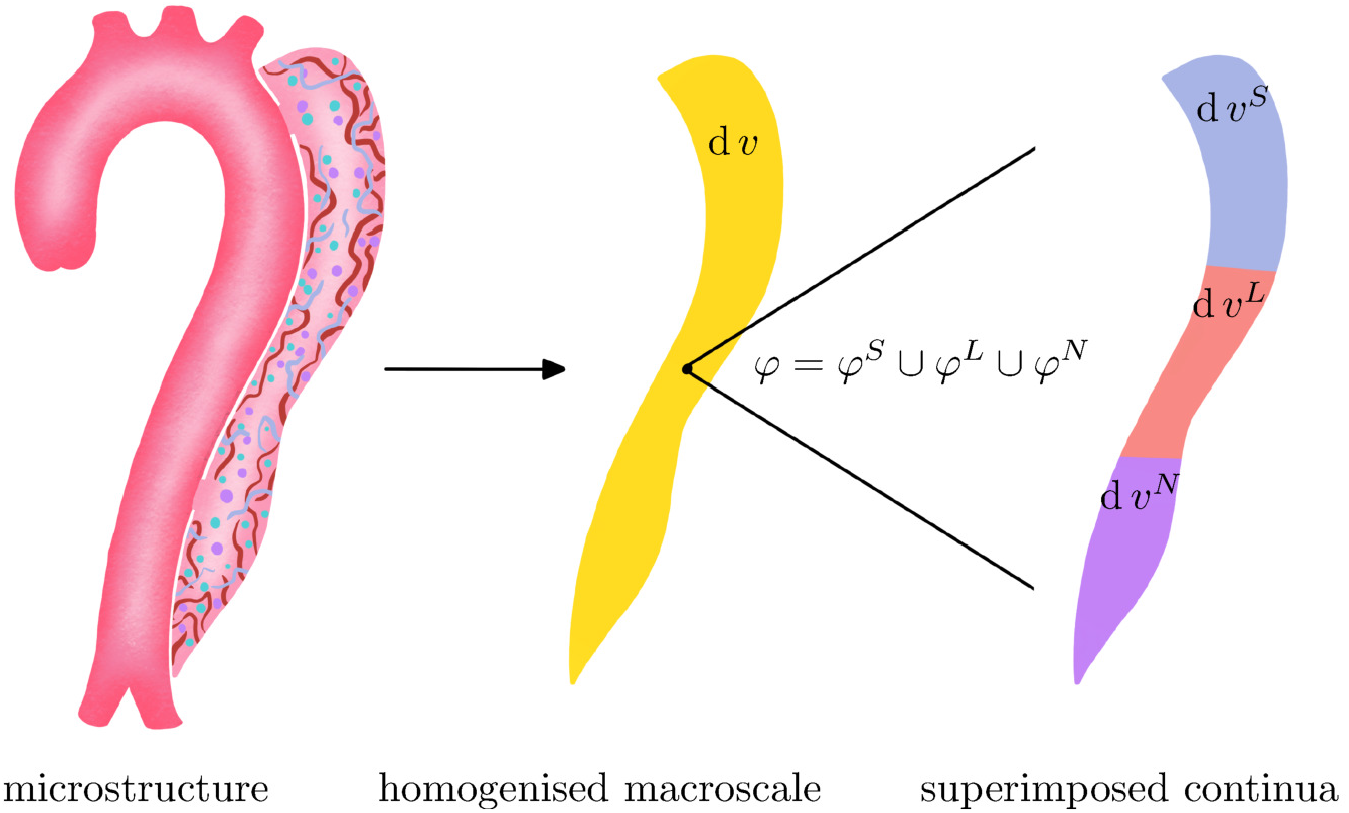
Illustration of the microstructure of the false lumen (left), macro-model obtained by volumetric homogenisation (centre), and superimposed continua (right)

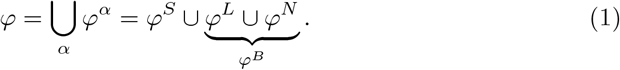

These three phases comprise the solid *φ*^*S*^, the liquid *φ*^*L*^, and the nutrient *φ*^*N*^. The solid phase *φ*^*S*^ consists of subendothelial collagen, activated platelets (which move to the injury site), fibrin fibres (formed during the coagulation cascade), and wall cells. The solid phase is saturated with fluid blood *φ*^*B*^. The blood itself consists of the nutrient phase *φ*^*N*^, containing deactivated platelets (activated upon injury) and clotting factors (part of the coagulation cascade), and the liquid phase *φ*^*L*^, representing blood excluding the nutrients. For further insights into TPM, the reader is referred to Ehlers (2002).

The concept of volume fractions *n*^*α*^ allows us to characterise the microstructure. It is defined as a ratio of the respective partial volume element d *v*^*α*^ to the bulk volume element d *v* of the multiphasic aggregate

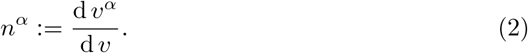

Furthermore, the partial density *ρ*^*α*^ of a constituent is defined as the ratio of the mass element of the constituent d *m*^*α*^ to the bulk volume element d *v*. Similarly, the realistic density *ρ*^*αR*^ of a constituent is the ratio of the mass element of the constituent d *m*^*α*^ to the constituent’s volume element d *v*^*α*^

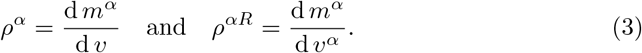

By substituting equation (2) into (3), we obtain the relation between the densities as

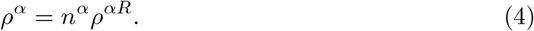

This implies that the material’s incompressibility (*ρ*^*αR*^ = constant) does not guarantee bulk incompressibility (*ρ*^*α*^ = constant) because the volume fraction can change owing to the deformation or mass exchange between phases.

### 2.2 Kinematics

In the context of the superimposed and interacting porous continua, particles *P*^*α*^ of all the constituents *φ*^*α*^ simultaneously occupy each spatial point **x** of the current configuration at any time *t*, cf. Figure 3. Since the particles at **x** proceed from the different reference positions **X**_*α*_ at time *t*_0_, each constituent is assigned its own motion and velocity functions

**Fig. 3.**
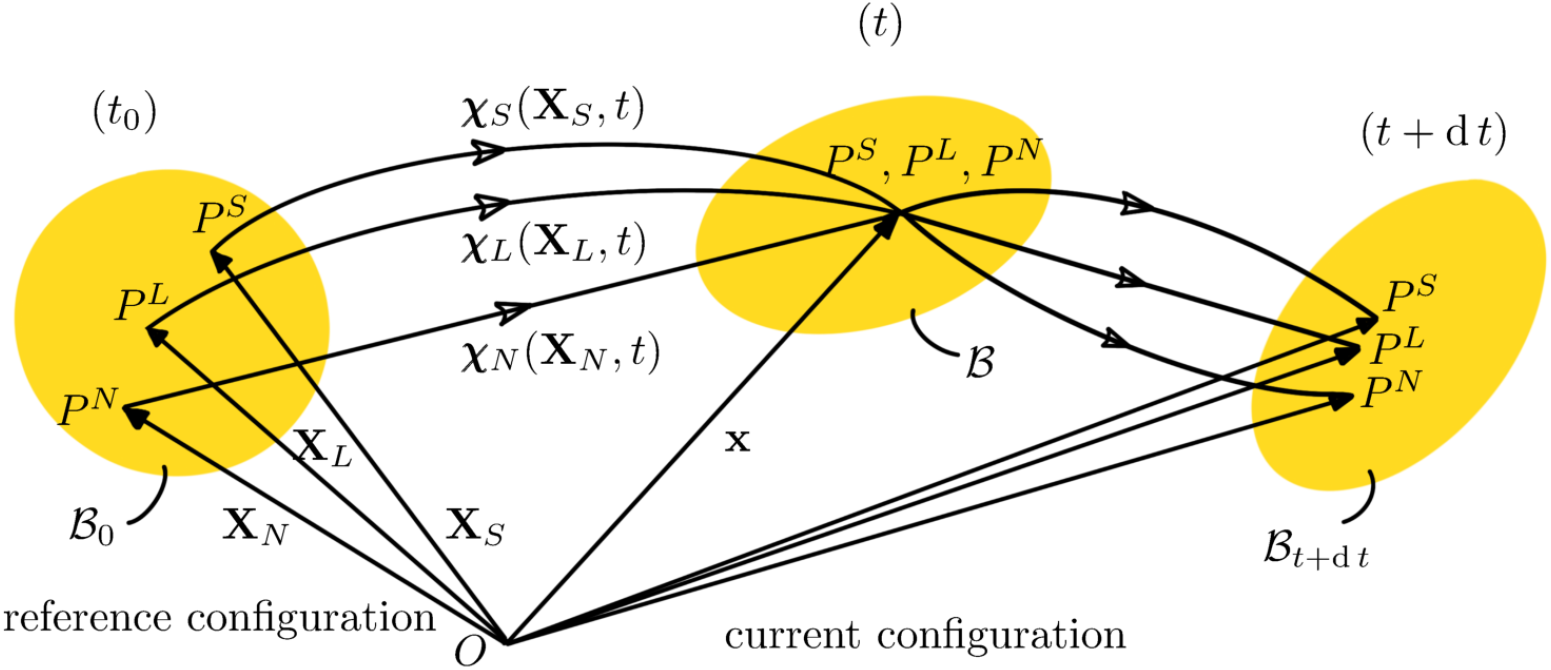
Motion of a triphasic aggregate (Gupta, 2023)

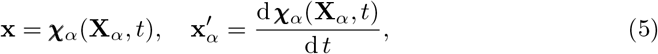

where 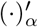 is a material time derivative defined as 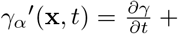 grad 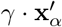 for an arbitrary scalar-valued field function *γ*_*α*_. A unique inverse motion function 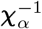 must exist to have a unique reference position **X**_*α*_ at time *t*_0_ for each material point *P*^*α*^ at **x**. The necessary and sufficient condition for this is the existence of a non-singular Jacobian *J*_*α*_

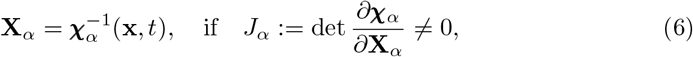

where det(·) denotes the determinant operator. Furthermore, the deformation gradient and its inverse can be defined as

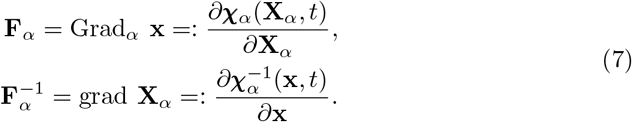

The right Cauchy-Green deformation tensor **C**_*α*_ is defined as 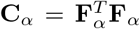 and the left Cauchy-Green tensor **B**_*α*_ is defined as 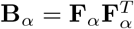. Moreover, the spatial velocity gradient 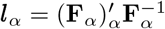 can be additively decomposed into the symmetric and skew-symmetric parts

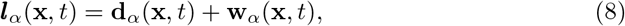

Where

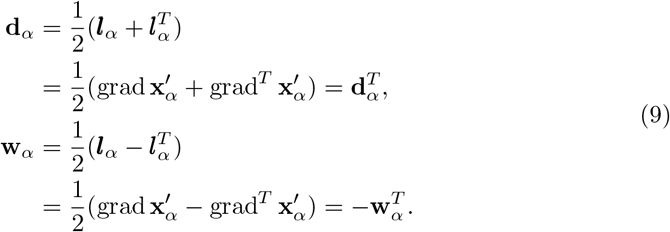

**d**_*α*_ is the symmetric part and is referred to as the rate of deformation tensor (rate of strain tensor), and the skew-symmetric part **w**_*α*_ is known as the spin tensor (rate of rotation tensor or vorticity tensor).

### 2.3 Balance Relations

After a brief introduction to the TPM and the kinematic quantities, we introduce the local balance relations for the constituents *φ*^*α*^. The mass balance is given by

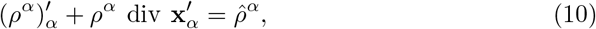

where 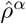 is the mass production term of the constituent *φ*^*α*^. The volume balance can be obtained using the above equation and the equation (2) as

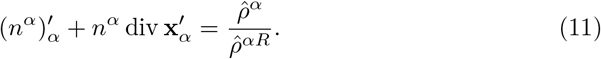

The balance of momentum for quasi-static conditions is read as

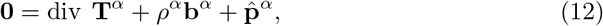

where **T**^*α*^ is the partial Cauchy stress tensor and 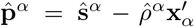 is the direct momentum production term. 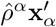 represents the momentum production due to the mass exchange and ŝ^*α*^ is the total momentum production. The moment of momentum balance leads to symmetric stress tensors

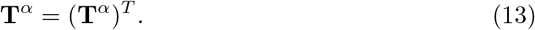

The total production terms are restricted by the summation constraints

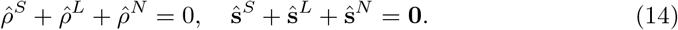

### 2.4 Assumptions

Thereafter, we present meaningful assumptions to simplify the model to a reasonable scope and adapt the model according to the theoretical background of thrombosis.

- The aggregate is assumed to be fully saturated. This condition has to be permanently fulfilled to avoid the formation of any vacant space

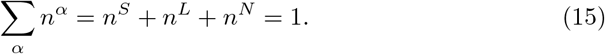
- The constituents are assumed to be materially incompressible but it does not lead to bulk incompressibility

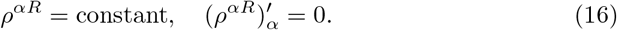
- Isothermal conditions are assumed. Therefore, it is not needed to use the energy balance explicitly

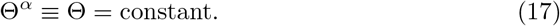
- Body forces are neglected

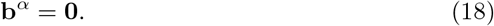
- Both the nutrient and the liquid phases are a part of the blood phase. Therefore, both phases are assumed to have the same velocity

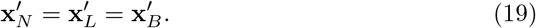
- Furthermore, the mass exchange occurs between the nutrient and the solid phases from the physiological understanding. The liquid phase is not involved in the mass transfer

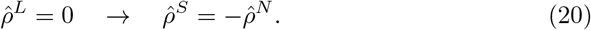

Additionally, the internal structure of a thrombus is considered to be isotropic, quasistatic conditions are assumed, and the diffusion velocity is neglected.

### 2.5 Constitutive Modelling

Thereafter, we follow the principles of determinism, equipresence, local action, and material frame indifference. Further information regarding these principles specific to this application can be found in the thesis by Gupta (2023). Moreover, to restrict the complexity of the model to a reasonable scope, the dependencies of the Helmholtz free energy are chosen as

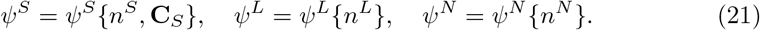

It is essential to recognise that the solid volume fraction *n*^*S*^ is dependent on det **F** and 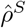 in the cases of incompressible constituents with mass exchange. This results from the time integration of the solid volume balance. However, we choose *n*^*S*^ as an independent constitutive variable to provide an additional degree of freedom for analysing the growth process. With *n*^*S*^ as the primary unknown variable, the saturation condition (15) also leads to *n*^*N*^ as an unknown variable. The splitting of *n*^*B*^ into *n*^*L*^ and *n*^*N*^ allows us to differentiate between the nutrients (deactivated platelets and clotting factors) and the liquid phase (blood excluding the nutrients) despite the velocity assumption (19). This helps to use the physiological understanding for modelling that the mass exchange occurs between the solid and the nutrient phases without altering the fluid content. The weak formulations of the governing equations for unknowns *n*^*S*^ and *n*^*N*^ will be presented in section 2.6.

Furthermore, the invariance condition and the polyconvexity are satisfied by representing the solid Helmholtz free energy function as

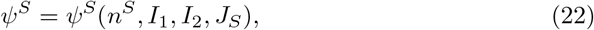

where *I*_1_ and *I*_2_ are principal invariants of **C**_*S*_. For a detailed discussion on convexity conditions, the reader is referred to Dacorogna (2007). Thereafter, the Helmholtz free energy function can be constructed as a modified Neo-Hookean material (Ricken et al. 2007; Gupta and Schanz 2023a; Gupta 2023)

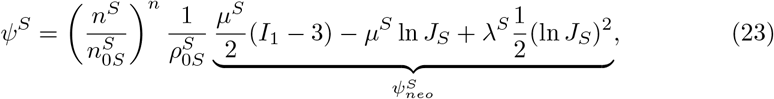

where 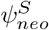 is the Neo-Hookean material law. *µ*^*S*^ and *λ*^*S*^ are the macroscopic Lamé constants. *I*_1_ = tr **C**_*S*_ is the first principal invariant of **C**_*S*_. (*·*)_0*S*_ represents the initial value concerning the reference configuration of the solid at time *t*_0_. The ratio of the volume fractions in the front accounts for the change in solid rigidity corresponding to changes in solid volume fractions. This helps to adapt the solid properties as the porosity changes. The material parameter *n* was identified to be 3 by Carter and Hayes (1977) and has been used in multiple TPM-based growth models (Ricken et al. 2007; Ricken and Bluhm 2009; Ricken and Bluhm 2010; Gupta and Schanz 2023a).

The earlier sections lead to a reduction in process variables. This simplifies the evaluation of the entropy inequality to

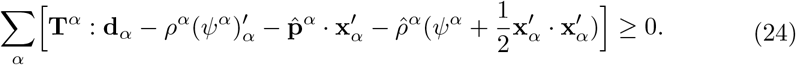

The detailed process of the entropy inequality can be found in our previous work (Gupta and Schanz, 2023a; Gupta, 2023). However, it’s important to note that in this study, we specifically consider the dissipation effects of fluids. As a result, the restrictions obtained from evaluating the entropy inequality and the constitutive relations are presented in the following sections.

#### Stress Equations

We follow the procedure outlined by Coleman and Noll (1974) to evaluate the entropy inequality and incorporate considerations of fluid frictional stresses. This yields the following restrictions for stresses

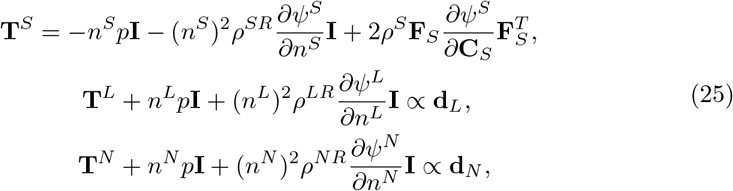

where signifies proportional relation, 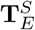 is the effective solid Cauchy stress and the Lagrangean multiplier *p* is identified as an unspecified pore pressure. Furthermore, we assume the blood to be a Newtonian and a homogeneous fluid. Therefore, considering *φ*^*B*^ = *φ*^*L*^ + *φ*^*N*^ with *∂ψ*^*B*^*/∂n*^*B*^ = 0 yields

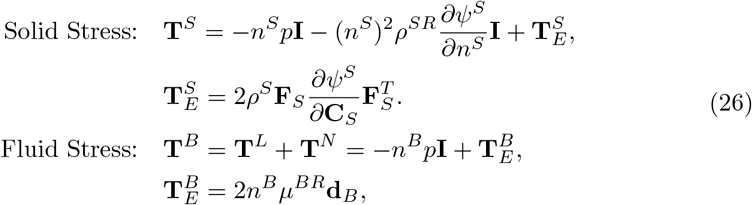

where **I** is the second-order identity tensor, 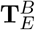 is the effective blood stress, and *µ*^*BR*^ [*N/m*^3^] is the dynamic viscosity of blood. Here, the fluid stress **T**^*B*^ is described as a sum of a pressure term *− n*^*B*^*p***I** and the deviatoric stress 2*n*^*B*^*µ*^*BR*^**d**_*B*_. Utilising equations (23) and (26), the relations for the effective solid Cauchy stress 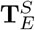 and the total solid Cauchy stress **T**^*S*^ can be expressed as

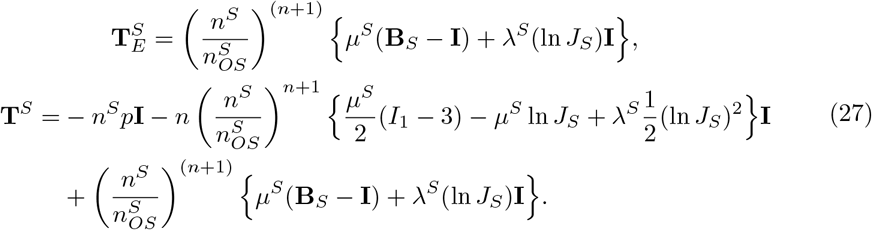

This gives the total Cauchy stress **T** as

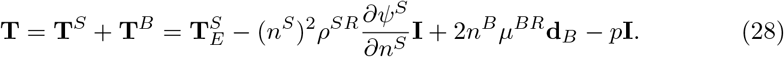

The blood velocity 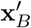 can be defined in relation to the solid velocity 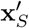 using the seepage velocity 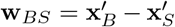. This seepage velocity characterises the fluid motion relative to the solid. The restriction obtained from the entropy inequality for the momentum production term 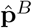 can be expressed in terms of the seepage velocity for an isotropic material as

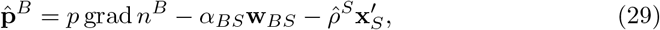

where *α*_*BS*_ can be described by using initial intrinsic permeability of solid 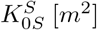 and dynamic blood viscosity *µ*^*BR*^ [*Ns/m*^2^] as

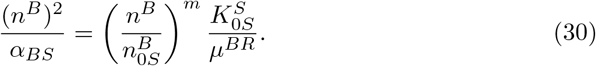

Thereafter, inserting the equations (29) and (30) in the fluid momentum balance (12), we obtain the Navier Stokes equation for quasi-static conditions extended to the porous media as

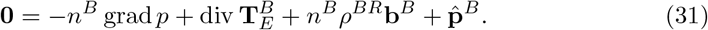

Using the equations (9)_1_, (26)_2_, and (31), the filter velocity *n*^*B*^**w**_*BS*_ can be formulated as

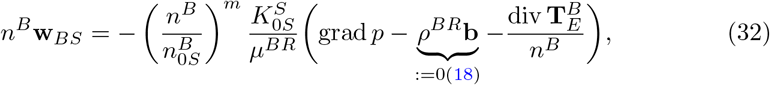

where

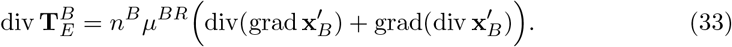

Here, 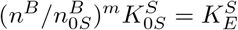 is termed as the effective permeability which considers the change of permeability due to evolving porosity. The porosity-dependent permeability was introduced by Eipper (1998), where *m ≥* 0 is a material parameter. 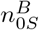 is the initial volume fraction of blood and *ρ*^*BR*^ is the realistic density of blood. Furthermore, equations (32) and (33) result in a modified version of a non-linear Darcy-Brinkman type equation

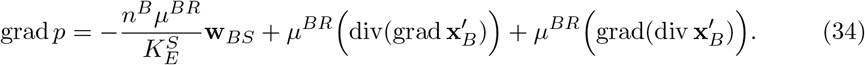

It is important to highlight that when considering a non-deforming solid 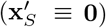 and no mass exchange 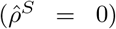, the aforementioned equation reduces to the Darcy-Brinkman equation. Ehlers (2022) has extensively reviewed the classical hydromechanical equations within the framework of TPM, providing valuable insights into their application and significance.

#### Mass Production

Based on the assumptions outlined in subsection 2.4, the mass exchange takes place between the solid and the nutrient phases, characterised by 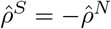. Subsequently, in accordance with the constraints derived from the entropy inequality, the chemical potential functions 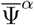 governing the solid growth process are related as

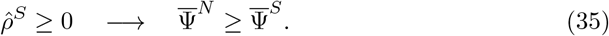

In consideration of the thrombus growth, we formulate the mass production based on the physiological understanding of the process. Virchow’s triad describes three physiological factors that influence the process of thrombosis. These factors are endothelial injury, hypercoagulability of blood and stasis of blood flow (Kumar et al. 2010). Endothelial injury refers to a vessel wall injury and triggers the platelet activation and coagulation process. Hypercoagulability means an increased tendency of coagulation in the body, and stasis of the blood flow refers to the condition of slow blood flow. These factors are considered extremely important to predict the growth of a thrombus (Stone et al. 2017). Motivated by Virchow’s triad, we dedicate our efforts to include all three factors in this model. An injury is already represented in the model through a false lumen. Also, the effects of the blood velocity and the nutrients on the thrombus growth are well researched and can be used to account for hypercoagulability and blood stasis (Zucker 1989; Begent and Born 1970; Pivkin et al. 2006). Therefore, the solid mass production 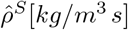 is postulated as a function of seepage velocity **w**_*BS*_ and nutrient volume fraction *n*^*N*^. Furthermore, it is understood that an initial porous thrombus forms rapidly at the injury site. The rate of the thrombus growth decreases as the solid thrombus increases. This decrease primarily occurs due to the reduced permeability, the restoration of normal blood velocity, and the decrease in blood stasis. The secondary haemostasis process continues for an extended duration, resulting in the gradual transformation into a solid thrombus over time. Thus, the solid mass production 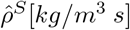 is postulated also as a function of the solid volume fraction. This leads to

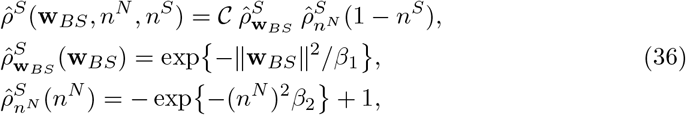

where *𝒞* represents the maximum mass exchange. 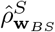 denotes the increased solid mass production during low fluid velocities, reflecting the stasis of blood. The material parameter *β*_1_ determines the dependence of mass exchange on fluid velocity, cf. Figure 4. Furthermore, 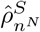 signifies the solid mass production in the presence of nutrients. The material parameter *β*_2_ can be used to adapt the model to various hypercoagulability scenarios. Additionally, a linear relation for the solid volume fraction *n*^*S*^ is added to the formulation to regulate the growth rate with an increase in solid volume fraction *n*^*S*^. However, obtaining data for the parameters in this approach can be challenging. Therefore, we opt for parameters that yield reasonable results.

**Fig. 4.**
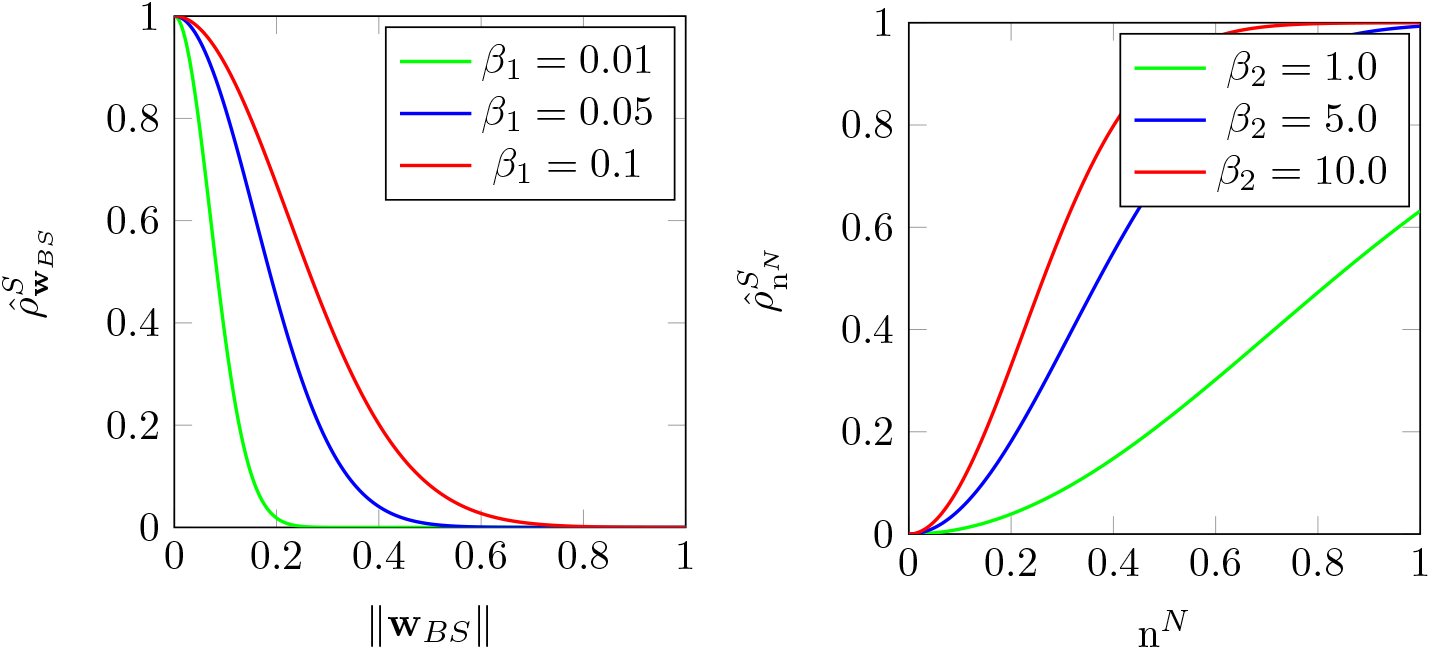
Variation in mass exchange rate dependence on the seepage velocity **w**_*BS*_ and nutrient volume fraction *n*^*N*^ influenced by the values of *β*_1_ and *β*_2_ (Gupta and Schanz, 2023a)

### 2.6 Numerical Treatment

Considering the assumptions, balance equations, and constitutive relations discussed in the preceding sections, we arrive at a set of five unknown variables:

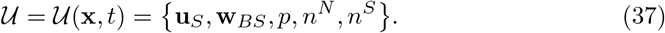

The next step is to formulate the weak formulation of the governing PDEs in the framework of the standard Galerkin procedure (Bubnov-Galerkin). Therefore, we introduce the weak forms of the momentum balance of the mixture, the momentum balance of the blood, the volume balance of the mixture, the volume balance of the solid, and the volume balance of the nutrients by multiplying them with respective test functions. Thereafter, we integrate the equations over the spatial domain Ω, occupied by the body *ℬ* at time *t*. We use the integration by parts and the divergence theorem to introduce the Neumann (natural) boundary terms. This yields a weak formulation of the governing equations.

- Momentum balance of mixture:

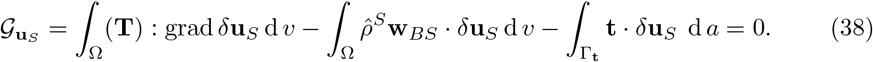
- Momentum Balance of blood:

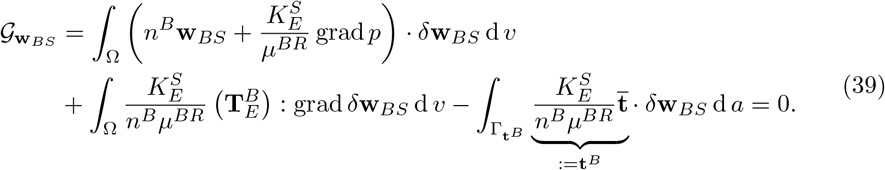
- Volume balance of mixture:

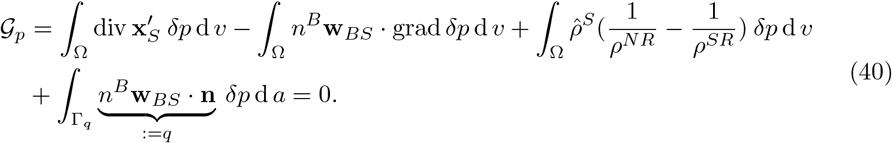
- Volume balance of nutrients:

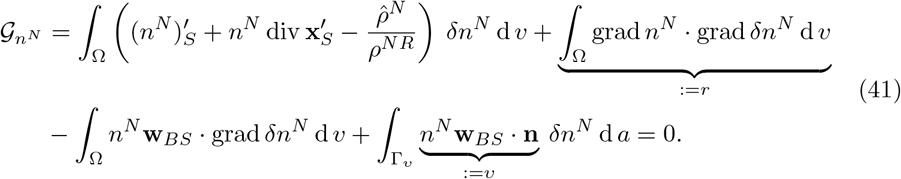
- Volume balance of solid:

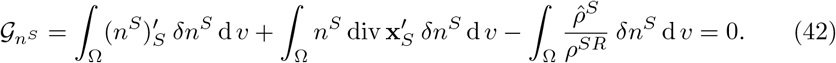

In the above set of weak formulations, **t** is the external load vector on the Neumann boundary Γ_**t**_, **t**^*B*^ is the external fluid load vector on the Neumann boundary 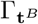, *q* is the fluid mass efflux on the Neumann boundary Γ_*q*_, and *υ* is the nutrient mass efflux on the Neumann boundary Γ_*υ*_, where **n** is the outward-oriented unit surface normal. Furthermore, the strong form of volume balance of nutrients is given by

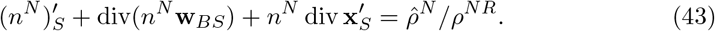

This equation has a typical structure of an advection equation with a source term on the right side. It is well-recognised that the weak form of such an equation tends to oscillate significantly unless stabilised or the mesh size is extremely small. To stabilise this equation, we choose the artificial diffusion scheme and add the term *r* to the equation (41). The reader is referred to Santos et al. (2021) and the references therein for further details on mass transport equations and stabilisation schemes.

Furthermore, we need a mixed formulation for the spatial discretisation because the unknowns have to be approximated simultaneously. Therefore, we choose quadratic shape functions for the solid displacement **u**_*S*_, and linear shape functions for the pressure *p*, the seepage velocity **w**_*BS*_, the solid volume fraction *n*^*S*^, and the nutrient volume fraction *n*^*N*^. This combination of the shape functions leads to the mixed formulation known as Taylor-Hood elements (Taylor and Hood, 1973). We use the same basis functions for the geometry as for the solid displacements **u**_*S*_ leading to the isoparametric mapping with respect to the solid displacements. Moreover, we use an implicit Euler scheme based on the backward Taylor-series expansion for the time discretisation. This gives a nonlinear system of equations which is solved using the Newton-Raphson scheme.

### 2.7 Geometry Preperation and Model Configuration

The weak formulation of the triphasic model is discussed in the preceding chapter. Additionally, the constitutive relations and assumptions establish the remaining thrombosis-specific relations. Next, we will describe the geometry.

Initially, we acquired the geometry of the type B AD of a 25-year-old female patient from the Vascular Model Repository (Wilson et al. 2013), where the clinical collaborators have carefully selected, unidentified, and provided data for this repository. The left image in Figure 5 depicts the aorta with a false lumen in the case of type B AD. Upon rotating the geometry, we observe the true lumen as well, cf. Figure 5 (centre). As our focus is on modelling thrombosis in the false lumen and TPM is more suitable approach for modelling porous materials like a thrombus, we extract only the false lumen from the complete geometry. This is achieved by making a perpendicular cut to the entry and exit tears. This gives us only the false lumen required for the simulation, cf. Figure 5 (right).

**Fig. 5.**
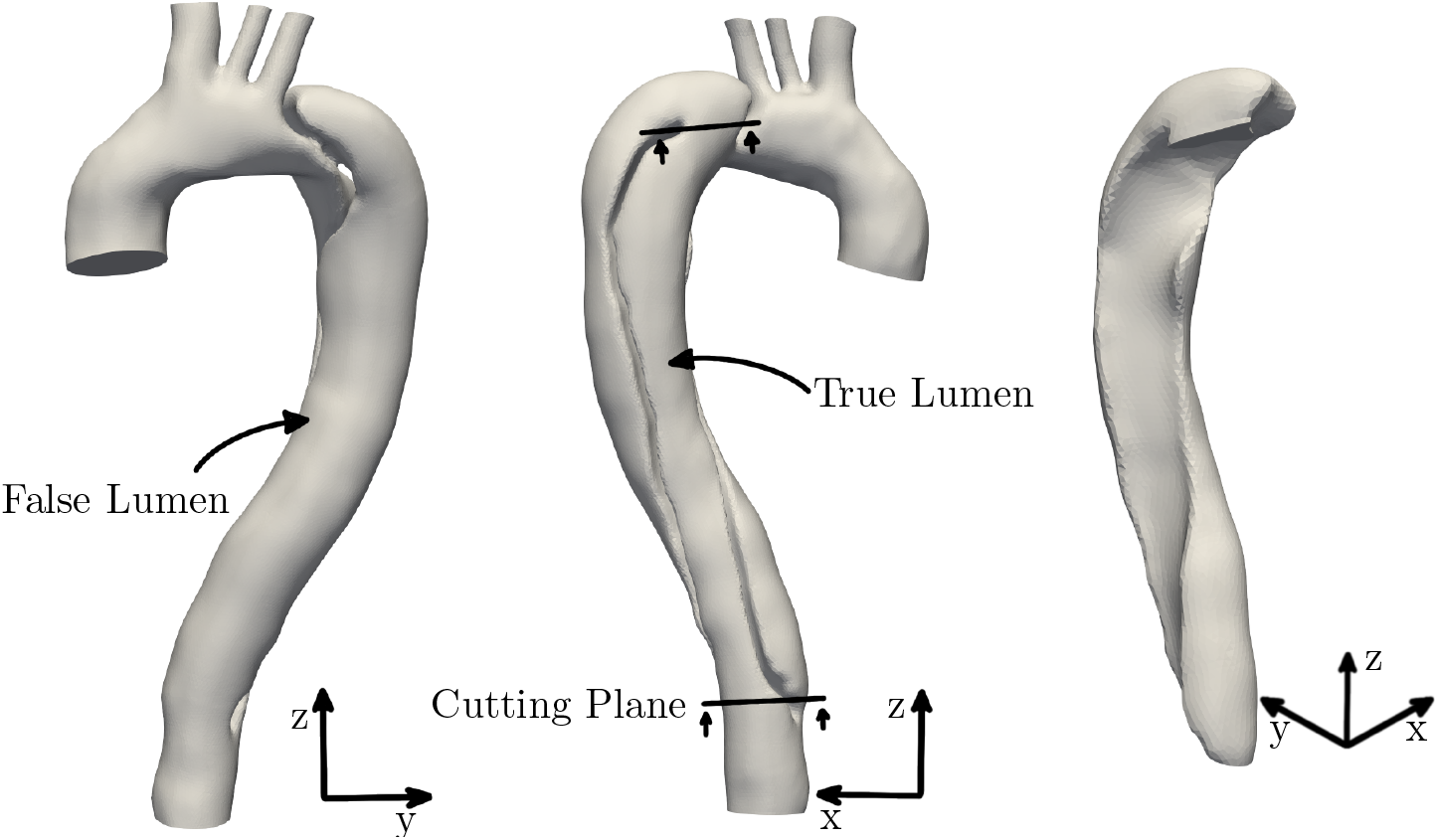
3D geometry of the aorta with type B AD showing the false lumen (left), true lumen and cutting planes (centre), and extracted false lumen (right)

This false lumen geometry represents a solid matrix saturated with the blood, comprising the liquid and the nutrient constituents. The entry tear at the top allows the blood to flow in. The Neumann boundary condition for the fluid volume efflux is set to *q* = 0.2 m/s, as shown in Figure 6. A recent study by Zimmermann et al. (2023) presents 4D-flow MRI data for the same geometry, showing that flow in the false lumen varies approximately between 0 ml/s and 100 ml/s over one cardiac cycle. However, in our study, we have neglected the shorter time scales associated with pulsating blood flow, such as systole and diastole, since thrombosis occurs over much larger time scales. Also, we aimed to prepare the geometry as closely as possible achieving the inlet area of 239 m^2^. To simplify the model and avoid complications related to different time scales, we used an average flow rate of 50 ml/s leading to inlet velocity of 0.2 m/s.

**Fig. 6.**
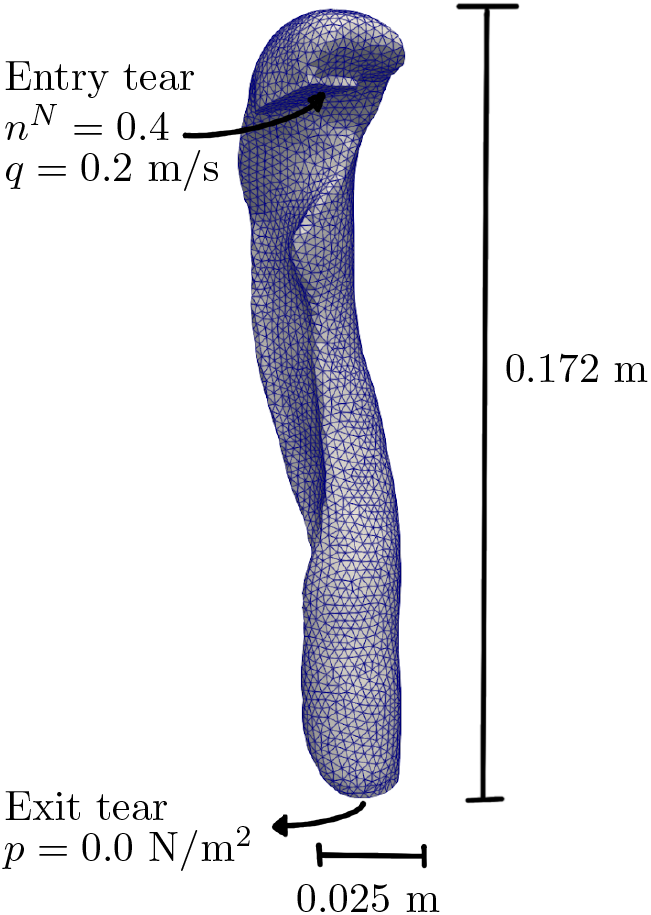
Discretisation and boundary conditions of the false lumen

Furthermore, a Dirichlet boundary condition is applied for the nutrient volume fraction, setting *n*^*N*^ = 0.4 at the entry tear. The exit tear is treated as a drained surface, with a pressure boundary condition set to *p* = 0.0 N/m^2^, allowing a free flow. Additionally, to promote thrombi growth at the walls, the seepage velocity **w**_*BS*_ is set to a small value of 0.01 m/s at the false lumen walls. In reality, the false lumen geometry is connected to the true lumen at the entry and exit tears, and to replicate this, we constrain the geometry at these tears in all directions. Furthermore, the simulation is performed using the parameters listed in Table 1, with a time step size of 100 seconds with a total time of 300000 seconds (*∼* 83.33 hours). For simplicity and due to a lack of material data, we assume a zero Poisson’s ratio (*λ*^*S*^ = 0 N/m^2^) for the solid skeleton (Zheng et al. 2020; Taylor et al. 2014; Yang et al. 2021). Furthermore, we implemented the model using FEniCSx, an open-source platform for solving partial differential equations with the finite element method. To handle the nonlinear aspects of the problem, we employed the Newton solver. We have set both absolute and relative tolerances to 1*×* 10^*−*8^ and used incremental convergence criteria, focusing on changes in the solution between iterations.

**Table 1.** Parameters for modelling thrombosis (Gupta and Schanz, 2023a)

| Parameter | Value | Unit | Parameter | Value | Unit |
| --- | --- | --- | --- | --- | --- |
| $\lambda^S$ | 0.0 | N/m<sup>2</sup> | $\rho^{SR}$ | $2 \times 10^3$ | kg/m<sup>3</sup> |
| $\mu^S$ | $1 \times 10^5$ | N/m<sup>2</sup> | $\rho^{LR}$ | $1 \times 10^3$ | kg/m<sup>3</sup> |
| $\mu^{BR}$ | $1 \times 10^{-3}$ | Ns/m<sup>2</sup> | $\rho^{NR}$ | $2 \times 10^3$ | kg/m<sup>3</sup> |
| $K_{OS}^S$ | $1 \times 10^{-6}$ | m<sup>2</sup> | $n_{OS}^S$ | 0.2 | - |
| $\mathcal{C}$ | $5 \times 10^{-2}$ | kg/m<sup>3</sup>s | $n_{OS}^N$ | 0.4 | - |
| $\beta_1$ | 0.05 | s<sup>2</sup>/m<sup>2</sup> | $\beta_2$ | 5.0 | - |
| $m$ | 3.0 | - | | | |

Moreover, to analyse the quality of spatial discretisation, we discretise the 3D false lumen using Gmsh with approximately (a) 65000, (b) 37000 elements, (c) 23000 elements, and (d) 10500 elements, cf. Figure 7. We perform simulations for all four meshes using the specified boundary conditions and the parameter values given in Table 1. Thereafter, we plot the average value of the solid volume fraction over the whole domain and compare the results. In Figure 8 (left), we observe the solid volume fraction converges to values between 0.970 and 0.963. The zoomed-in plot reveals that the values for meshes (a) and (b) are further away compared to the meshes (c) and (d). The values of the solid volume fraction at the end time are 0.9637 and 0.9639 for the meshes (a) and (b) respectively. Based on these findings, we conclude that the solution converges more effectively for meshes (a) and (b). However, using mesh (a) with 65000 elements would significantly increase the system size without considerably improving results. Therefore, we choose the mesh (b) with 37000 elements for further simulations.

**Fig. 7.**
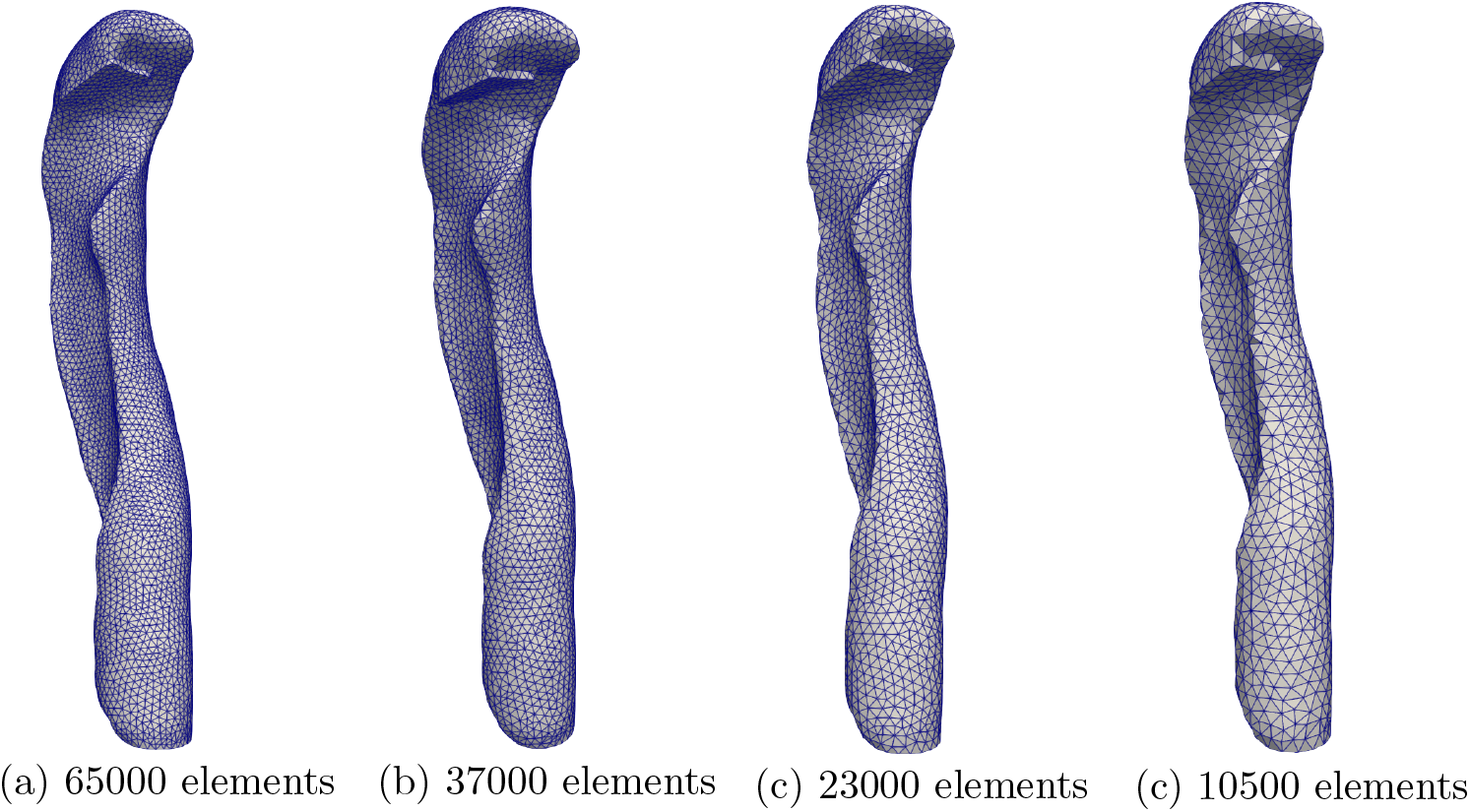
Discretisation of the geometry with different mesh sizes.

**Fig. 8.**
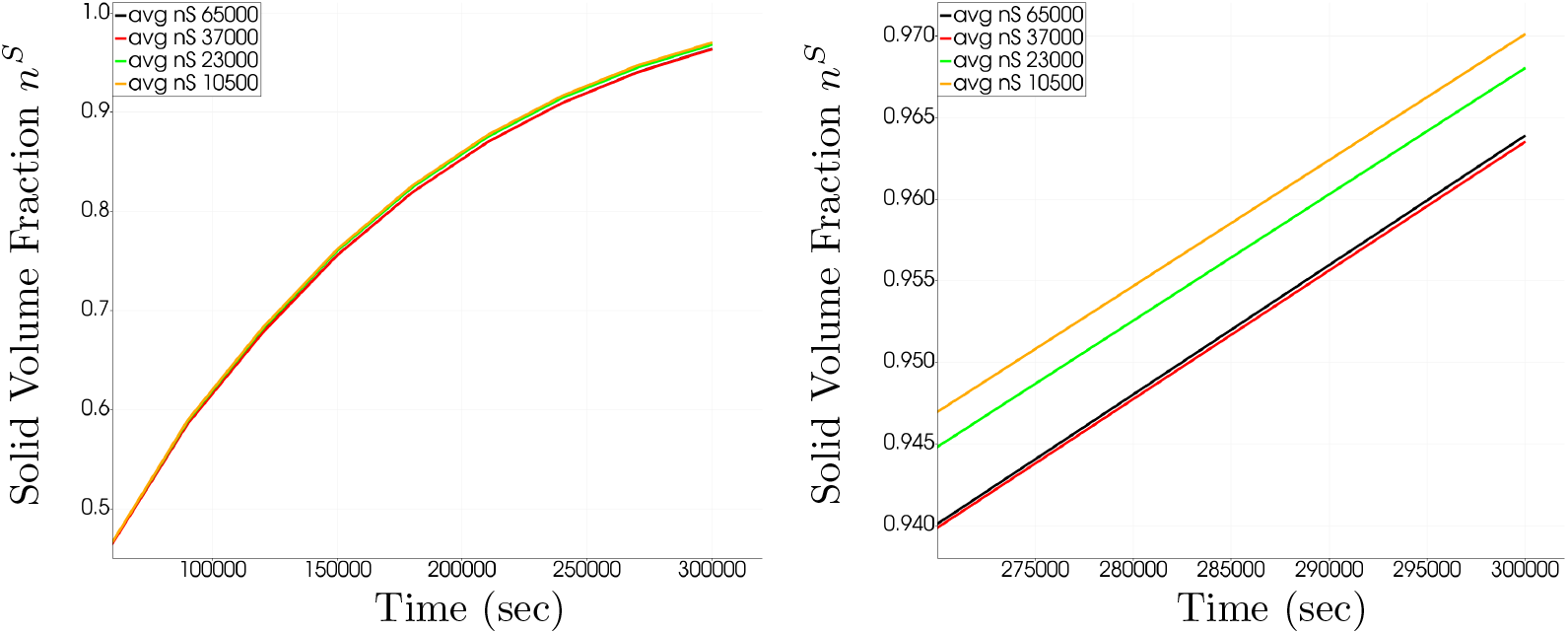
Convergence study of the average solid volume fraction over the whole domain with the overall (left) and zoomed-in (right) plots.

## 3 Results of Thrombosis Modelling in Type B Aortic Dissection

In this section, we demonstrate the application of the developed model for thrombosis using 3D real geometries for type B AD. Firstly, we define three planes (P1, P2, P3) passing through the false lumen to analyse the results, as illustrated in Figure 9 (left). Upon performing the simulation with the specified setup and parameter values, we obtain solid volume fraction plots at time *t* = 300000 seconds, as depicted in Figure 9 (centre). The cross-sectional views at pre-defined planes indicate the formation of the thrombus and the flow of blood in certain parts of the false lumen. The solid volume fraction plots in Figure 9 (right) show that the thrombus is formed on the walls of the lumen which are the sites of injury. These observations align with our physiological understanding of thrombosis. Moreover, since a thrombus in such cases develop over days or weeks, the total time of 300000 seconds is considered to demonstrate the capability of the model to predict thrombi growth within a realistic time frame. Furthermore, the model demonstrates its capability to represent blood pressure, a major risk factor for Aortic Dissection, which significantly impacts thrombi growth. To investigate this, we assign three different values to the fluid volume efflux *q*: (a) 0.08 m/s, 0.2 m/s, and (c) 0.4 m/s. The results of these three cases at plane P1 are shown in Figure 10. The varying values of the entry tear boundary condition *q* is capable to represent the range of blood pressures. A higher value of *q* corresponds to higher blood pressure, thereby making thrombi formation more challenging, and vice versa, aligning well with physiological expectations.

**Fig. 9.**
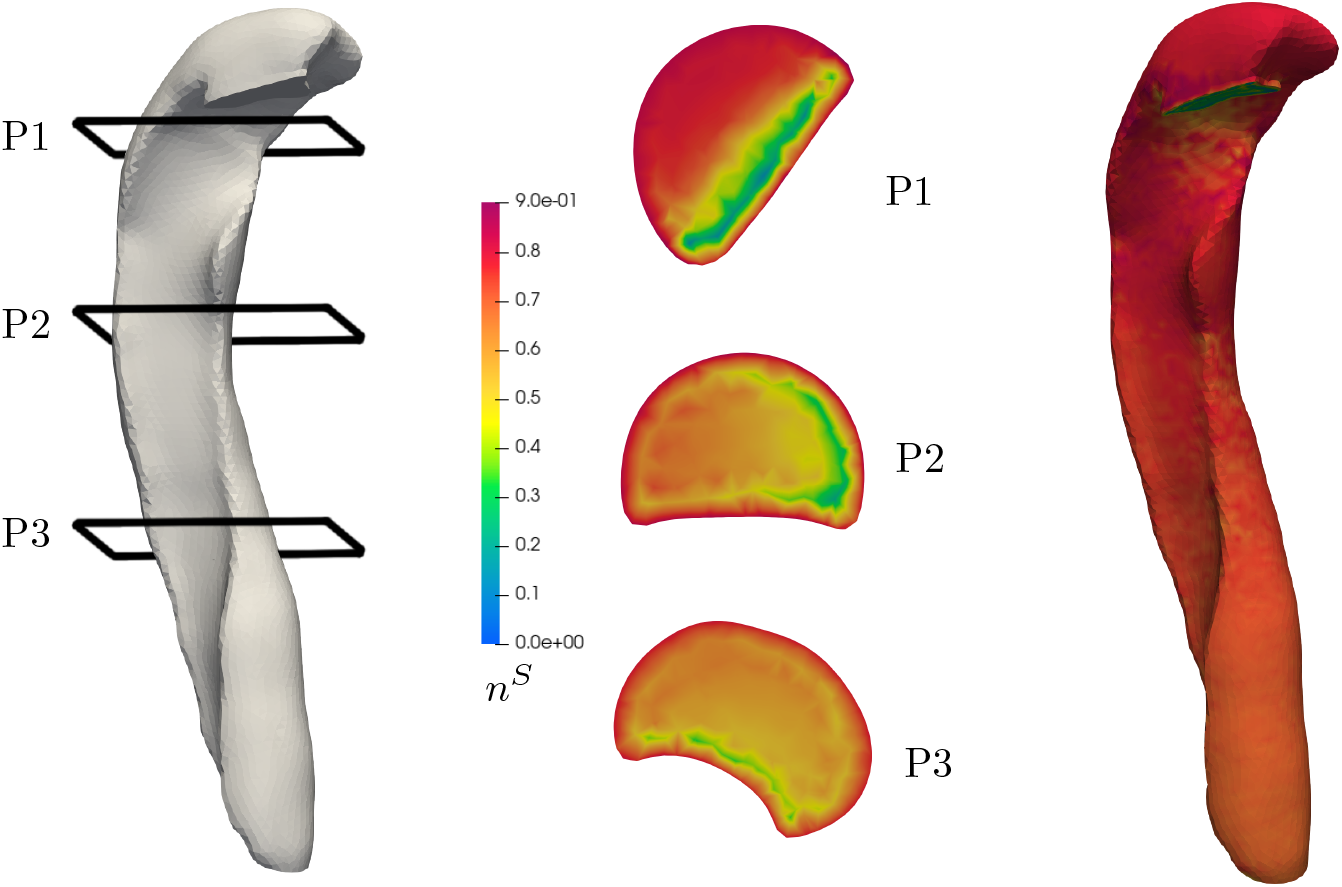
Geometry of the false lumen with three planes named P1, P2, and P3 (left), solid volume fraction plots illustrating the thrombus at the cross sections of the designated planes (center), and the visualisation of the thrombus formation along the walls of the false lumen (right) at time *t* = 300000 seconds

**Fig. 10.**
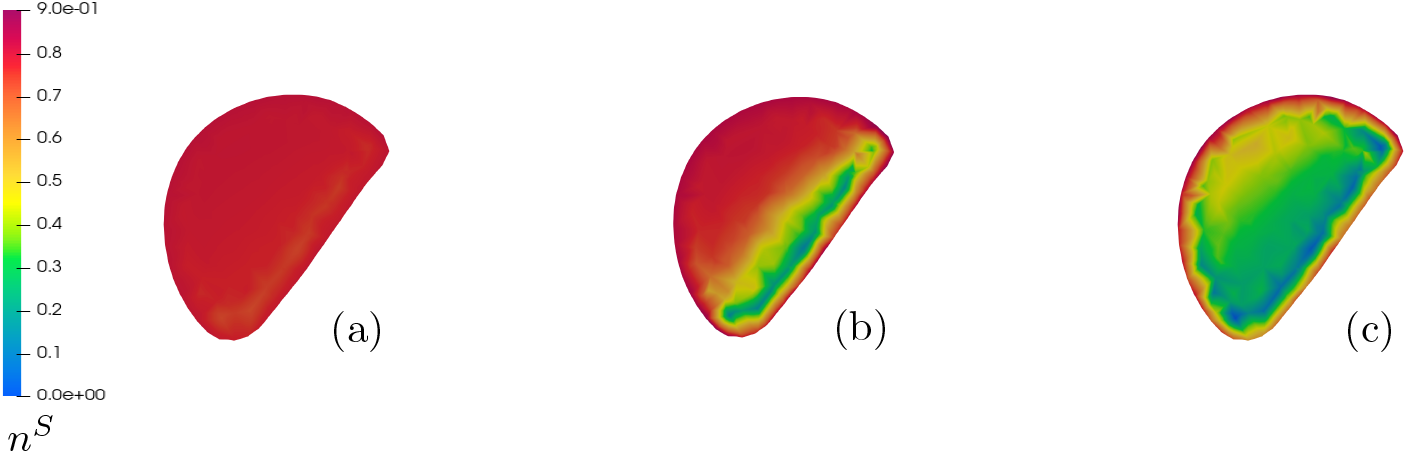
Visualisation of the thrombus at plane P1 demonstrates the impact of various values of the boundary condition *q* at the entry tear with *q* set to (a) 0.08 m/s, (b) 0.2 m/s, and (c) 0.4 m/s at time *t* = 300000 seconds

Furthermore, we can vary the values of parameters *β*_1_ and *β*_2_ in mass production equation (36). The different possible values of these parameters are visualised in Figure 4. The value of *β*_1_ affects the range of seepage velocity over which the thrombus grows. A lower value makes thrombus formation more difficult, while a higher value facilitates easier thrombus formation. Additionally, changing the values of *β*_2_ alters the relationship between mass production and nutrient volume fraction. Higher values of *β*_2_ promote easier thrombus formation, while lower values correspond to slower thrombus formation. These effects of *β*_1_ and *β*_2_ can be found in Gupta and Schanz (2023a).

However, validating thrombi growth is currently challenging due to the lack of data for the presented geometry of type B Aortic Dissection. Nonetheless, a study conducted by Zimmermann et al. (2023) provides 4D-flow MRI results for the geometry used in this investigation, offering an opportunity to validate the velocity results. To facilitate this comparison, we selected two additional geometries from the Vascular Model Repository, which were also used in this study. This brings the total number of geometries to three for comparison of the velocity results with the 4D-flow MRI data. We denote the original geometry as Case I. Case II features an exit tear area of approximately 25% of Case I, while Case III presents an entry tear area of approximately 25% of Case I. Figure 11 shows the entry and exit tear areas in all three cases. For the simulations, we assumed a constant inflow volume. Table 2 presents a quantitative comparison of velocity measurements obtained from the Theory of Porous Media (TPM) simulations at 1000 seconds and 4D-flow MRI data at peak systole across the three different false lumen configurations: Case I (Original), Case II (Smaller Exit), and Case III (Smaller Entry). Each configuration was analysed at three planes (P1, P2, P3), focusing on average velocities. TPM/MRI in the table represents the ratio of velocities from the TPM simulations to those measured by MRI.

**Table 2.** Quantitative comparison of TPM simulations and 4D-flow MRI velocities for different false lumen configurations.

| Planes | Case I: Original |  |  | Case II: Smaller Exit |  |  | Case III: Smaller Entry |  |  |
| --- | --- | --- | --- | --- | --- | --- | --- | --- | --- |
|  | Average [m/s] |  | TPM/MRI | Average [m/s] |  | TPM/MRI | Average [m/s] |  | TPM/MRI |
|  | TPM | MRI |  | TPM | MRI |  | TPM | MRI |  |
| P1 | 0.1509 | 0.2575 | 0.58 | 0.1466 | 0.0996 | 1.47 | 0.1539 | 0.0840 | 1.83 |
| P2 | 0.1581 | 0.2530 | 0.62 | 0.1539 | 0.0938 | 1.64 | 0.1605 | 0.0810 | 1.98 |
| P3 | 0.1423 | 0.2430 | 0.58 | 0.1386 | 0.0876 | 1.58 | 0.1451 | 0.0809 | 1.79 |

**Fig. 11.**
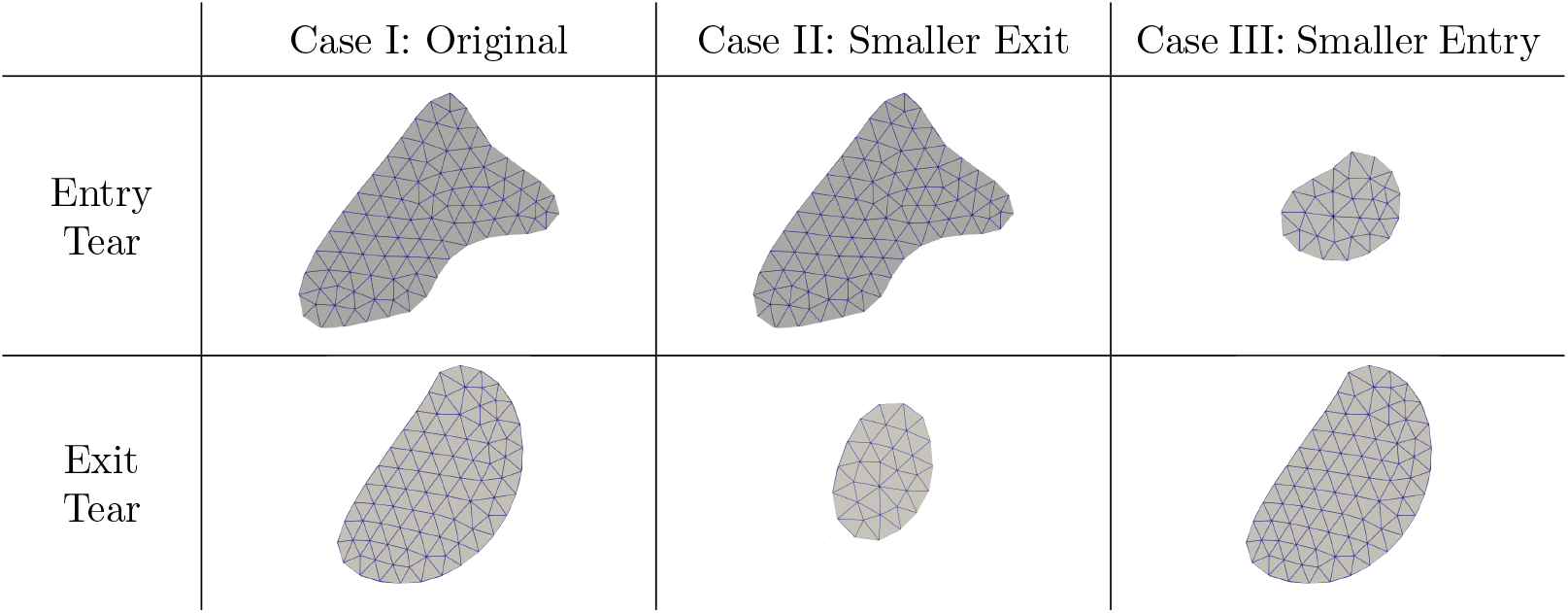
The 2D entry and exit tear areas for all three cases

**Fig. 12.**
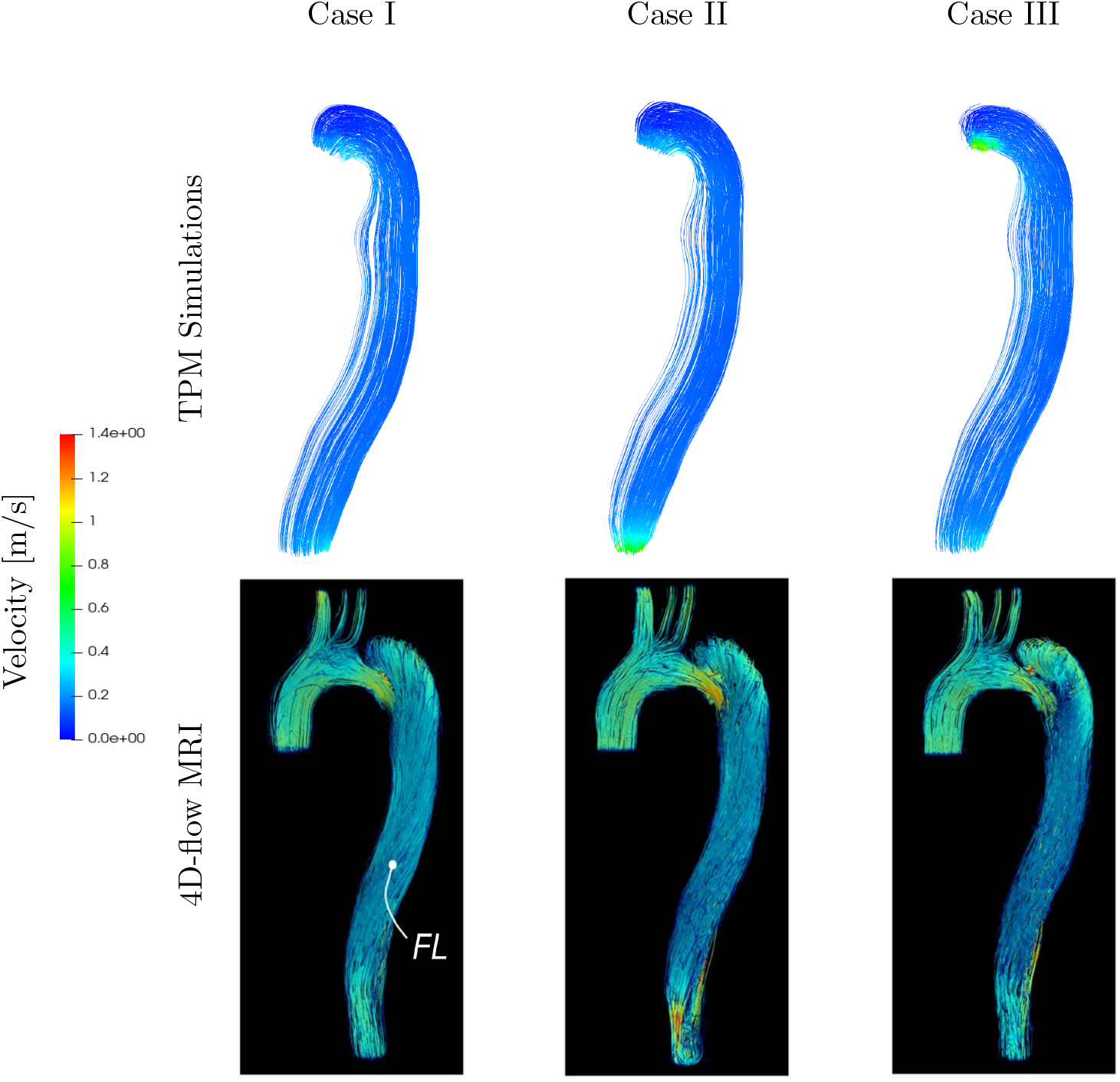
Comparison of velocity profiles: The top row displays the seepage velocity acquired from simulations employing the developed triphasic model. The bottom row shows the velocity obtained from the 4D-flow MRI with FL representing the false lumen (Adapted from Zimmermann et al. (2023) and used under [CC BY 4.0](https://creativecommons.org/licenses/by/4.0/))

For Case I, the TPM velocities are lower than the MRI values, with TPM/MRI ratios ranging from 0.58 to 0.62, indicating that the model underestimates flow compared to MRI. This trend reverses for the other two cases, with simulation velocities exceeding MRI values, resulting in ratios between 1.47 and 1.64 for Case II and between 1.79 and 1.98 for Case III. The data reveal distinct trends in how the TPM model aligns with the MRI measurements across these configurations. However, it should be noted that the MRI data provides velocity measurements over a single cardiac cycle, capturing fluctuations of blood flow at different stages. In contrast, the TPM simulations model flow and thrombus development over much longer timescales (days/weeks), focusing on the macroscopic evolution of fluid and solid interactions. Therefore, while direct comparisons with specific phases of the cardiac cycle are not feasible, the objective is to ensure that the model computes velocities within the expected physiological range. However, the observed discrepancies in results can be attributed to several factors. The assumption of constant flow in our simulation, which simplifies the complex, pulsatile nature of real blood flow. The model neglects shorter time scales, such as the systole and diastole phases, which could have a pronounced effect on the flow. Also, incorporating true lumen would influence flow and pressure dynamics further influencing flow distribution. Moreover, TPM simplifies the fluid flow by assuming a saturated matrix where blood and thrombi interact at a continuum scale. However, in a realistic scenario, the interaction between the blood flow and the walls of the false lumen, especially with thrombus growth, involves highly non-linear and complex behaviors, which may not be captured adequately in this approach.

Moreover, Figure 13 shows the evolution of the velocity field over time for Case I. It can be observed that the fluid velocity increases with time as more thrombi form, restricting the fluid flow as anticipated. Although this trend cannot be validated due to the lack of thrombosis data for this case, it provides valuable insight into how velocities in the false lumen may evolve as thrombi develop.

**Fig. 13.**
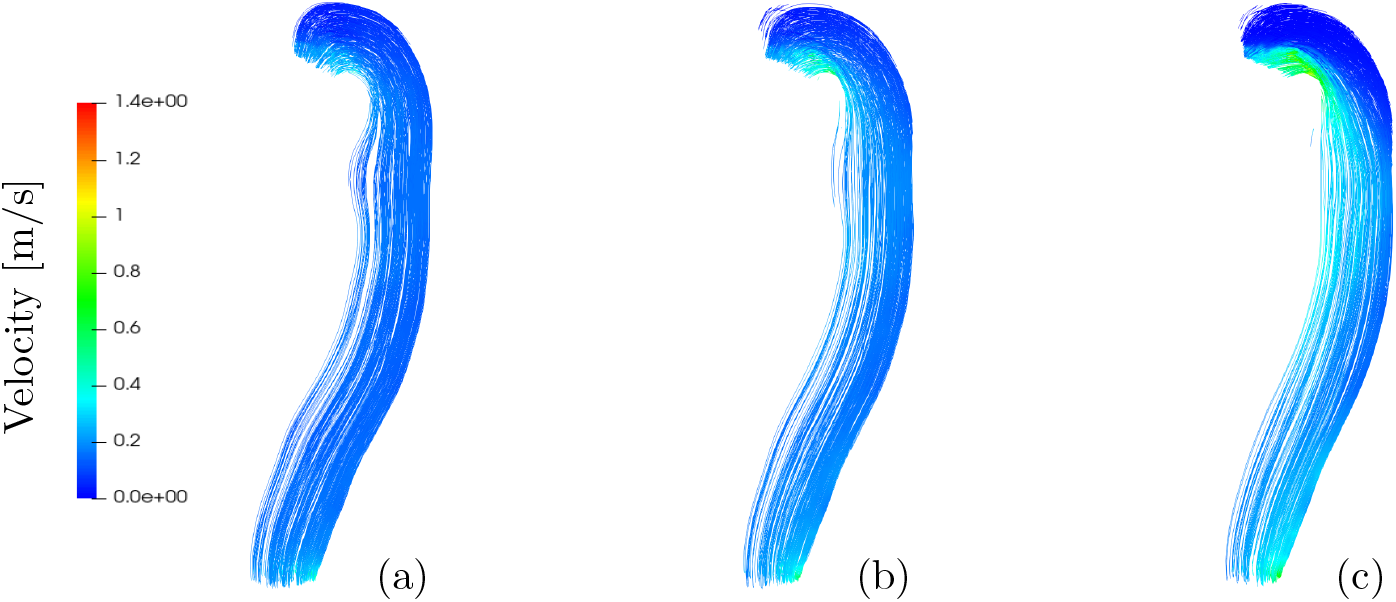
Evolution of seepage velocity at time *t* = (a) 1000 seconds, (b) 150000 seconds, and (c) 300000 seconds

## 4. Conclusion

The proposed triphasic model presents a promising approach to simulate thrombosis numerically. It offers the advantage of incorporating the mass exchange between the nutrient and the solid phases without altering the liquid volume, aligning well with the physiological understanding of thrombosis. Our analysis of the results reveals initial thrombi formation along the walls of the false lumen which is consistent with the process of platelet plug formation at injury sites. Over time, thrombi growth progressively restricts the blood flow through the cross-section of the false lumen. Moreover, we have integrated the representation of Virchow’s triad factors, alongside material parameters *β*_1_ and *β*_2_, enabling adaptation for various hypercoagulability and blood stasis scenarios. Additionally, the model can capture blood pressure, a significant risk factor in Aortic Dissection. The model demonstrates its utility in real-world scenarios such as type B AD while predicting thrombosis over a representative time scale. Moreover, the model yields velocities close to the MRI data. However, certain discrepancies highlight the need for further improvements of the model.

The growth process needs to be improved to capture all the essential features comprehensively. To fully model the thrombosis process, it is necessary to incorporate the chemical reactions involved in haemostasis and account for the non-Newtonian nature of blood. Additionally, assigning physical significance to the parameters is essential for the practical applications. Moreover, this study neglects the effects of pulsating blood flow, which occurs on shorter time scales, and instead considers a steady blood flow. However, it is important to note that pulsating blood flow may reduce stagnation in regions of low stress and affect thrombosis. Furthermore, only the false lumen geometry is considered in this study. We expect that incorporating the true lumen would impact flow and pressure dynamics. Specifically, backflow of blood during different phases of the cardiac cycle would influence flow redistribution between the false lumen and true lumen. Including the true lumen in the model is expected to provide a more comprehensive view and improve the accuracy of thrombosis predictions.

Although the presented model uses various simplifications and requires further refinement, it provides promising initial results and does not demand extensive resources. The development of biomechanical models presents challenges due to the lack of data and the biological complexity. Access to extensive medical and experimental data, including CT/MRI scans, is necessary for model validation, parameter determination, and refinement, leading to a more accurate representation of the growth process. Furthermore, the model’s extension to other medical conditions, such as arterial or venous thrombosis and DIC, holds the potential for revolutionising medical device design, personalised medicine, and disease prognosis. Understanding the mechanics of growth in chronic conditions could facilitate effective disease progression control and improve patient outcomes.

## Acknowledgement

We gratefully acknowledge Graz University of Technology for the financial support. Additionally, we acknowledge the Lead-project: Mechanics, Modeling, and Simulation of Aortic Dissection, as this research stems from the insights gained during this project.

## Statements and Declarations

- Competing interests: The authors have no relevant financial or non-financial interests to disclose.
- Data availability statement: The MSH files for the three false lumen geometries used in this research are available in the repository of Graz University of Technology (Gupta I, Schanz M (2024) False lumen geometries: Repository of msh files. https://doi.org/10.3217/cg2ch-ahe16).
- Ethical Statement: None

